# DNA methylation patterns of transcription factor binding regions characterize their functional and evolutionary contexts

**DOI:** 10.1101/2022.07.21.500978

**Authors:** Martina Rimoldi, Ning Wang, Jilin Zhang, Diego Villar, Duncan T. Odom, Jussi Taipale, Paul Flicek, Maša Roller

## Abstract

**Background:** DNA methylation is an important epigenetic modification which has numerous roles in modulating genome function. Its levels are spatially correlated across the genome, typically high in repressed regions but low in transcription factor (TF) binding sites and active regulatory regions. However, the mechanisms establishing genome-wide and TF binding site methylation patterns are still unclear.

**Results:** We used a comparative approach to investigate the association of DNA methylation to TF binding evolution in mammals. Specifically, we experimentally profiled DNA methylation and combined this with published occupancy profiles of five distinct TFs (CTCF, CEBPA, HNF4A, ONECUT1, FOXA1) in the liver of five mammalian species (human, macaque, mouse, rat, dog). TF binding sites were lowly methylated, but they often also had intermediate methylation levels. Employing a classification and clustering approach, we extracted distinct and species conserved patterns of DNA methylation levels at TF bound regions. CEBPA, HNF4A, ONECUT1 and FOXA1 shared the same methylation patterns, while CTCF’s differed. These patterns characterize alternative functions and chromatin landscapes of TF bound regions. Leveraging our phylogenetic framework, we found DNA methylation gain upon evolutionary loss of TF occupancy, indicating coordinated evolution. Furthermore, each methylation pattern has its own evolutionary trajectory reflecting its genomic contexts.

**Conclusions:** Our epigenomic analyses found that specific DNA methylation profiles characterize TF binding, and are associated to their regulatory activity, chromatin contexts, and evolutionary trajectories.

## BACKGROUND

Gene regulation is a complex process that controls gene expression across cell types and time points. Key players in establishing tissue-specific expression are transcription factors, which bind to specific DNA sequences, and covalent modifications to the DNA such as DNA methylation (DNAm). Regulatory evolution has widely been studied in comparative analysis of transcription factor binding, but complementary studies of the evolution of DNAm are lacking.

There are several known mechanisms influencing transcription factor binding evolution. Transcription factor (TF) binding evolves rapidly: in mammals it is characterized by frequent gain and loss of binding events even across short evolutionary time [1] [2][3] [4] [5]. Sequence divergence partly explains binding divergence. For example, a comparative study of a handful of liver-specific transcription factors in five mammals reported that more than 60% of binding losses could be explained by binding motif disruption through mutations or indels [6]. However, in the remaining 20-40% of lost binding events the motif was unchanged. Furthermore, TF binding in liver of rat and five mouse strains showed similar mutational rates between binding-conserved and binding-lost TF motifs [7], indicating that sequence divergence alone cannot explain TF turnover.

Despite the rapid rearrangements of the TF binding network [3][8], gene expression of orthologous genes tends to be conserved in mammals [9][10], likely due to the plasticity of the regulatory network [11]. For example, compensatory binding turnover in the proximity of lost events preserve regulatory network connectivity [11][6] and complexity of regulatory landscapes [12]. Finally, cooperative binding of multiple TFs [7] and clustered binding of a single TF [13] are more evolutionary stable than lone binding events. Less is known about how epigenetics modifications of DNA evolve and affect the evolutionary dynamics of transcription factor binding.

DNA methylation (DNAm) is a chemical modification of DNA, most commonly the addition of a methyl group to the fifth position of cytosines (5-methylcytosine (5mC)), for those cytosines followed by a guanine (CpGs). The presence of CpG methylation can be measured in bulk tissues and cell types as a continuous frequency value comprised between 0 to 100% (or 0 to 1) through whole-genome bisulfite sequencing assays (WGBS) [14]. Most CpGs in mammalian genomes measure 0-10% and 70-80% methylation, indicating overall unmethylated and methylated nucleotides, respectively [15]. However, about one in ten CpGs have intermediate levels, i.e. between 10 and 70% methylation [16], reflecting either the cell-to-cell variability of the bulk samples or epigenomic and transcriptional heterogeneity [17] [18] [19].

DNAm is largely recognized as a repressive epigenetic mark which often displays spatially correlated patterns across the genome [20]. Active regulatory regions, specifically CpGs islands and active promoters and enhancers, are typically unmethylated [21] [22] [23]. Genomic regions with intermediate methylation (IM) levels were shown to be widespread and conserved across species [24]. They typically co-localize with distal regulatory elements and can be reshaped upon transcription factor binding [16]. In fact, DNAm levels are tightly linked to functional and chromatin contexts. Transcriptional activity, TF binding, and chromatin remodelers have an impact on passive and active enzymatic processes that ultimately determine local patterns of methylation across the genome [25]. As a consequence, DNAm levels are highly predictive of regulatory activity [26] [27] [28] [29].

DNA methylation was traditionally thought to inhibit transcription factor binding by physically preventing proteins from binding their target DNA sequences [30]. However, mounting evidence from *in vivo* and *in vitro* experiments now challenge this view. High-throughput *in vitro* assays such as protein microarrays and methyl-SELEX (systematic evolution of ligands by exponential enrichment) showed that TFs not only bind methylated motifs, but also that their binding affinity can be enhanced by 5mC [31] [32] [33]. Evidence that the methylation landscape can be remodeled *in vivo* by the binding of specific transcription factors such as the CCCTC-binding factor (CTCF) or the RE1 Silencing Transcription Factor (REST) has further challenged the traditional view [16]. Despite the experimental *in vivo* identification of a handful of TFs with modified specificity for methylated motifs [34] [33], the impact of cytosine methylation on the regulation of TF binding in distinct genomic contexts remains unclear.

We designed a comparative epigenomic study of DNA methylation patterns within TF binding regions. We generated whole-genome bisulfite-sequencing data from livers of five mammalian species (human, macaque, mouse, rat, and dog) and retrieved publicly available ChIP-sequencing data from five transcription factors in matched tissues. Four of the assayed transcription factors represent key components of the liver-specific regulatory network [35], namely: CCAAT/enhancer-binding protein alpha (CEBPA), hepatocyte nuclear factor 4 alpha (HNF4A), One Cut Homeobox 1 (ONECUT1, also known as HNF6), forkhead box protein A1 (FOXA1, also known as HNF3A). The final TF included in this study, CTCF, is a ubiquitous and multifunctional protein [36]. We used these datasets to characterize

DNA methylation at TF binding regions, find different DNAm patterns within distinct functional genomic elements, and show that DNA methylation and TF binding co-evolve.

## RESULTS

### Experimentally profiling DNA methylation in transcription factor binding regions

We obtained flash-frozen liver samples from five mammalian species (*Homo sapiens, Macaca mulatta, Mus musculus, Rattus norvegicus* and *Canis familiaris*) and performed whole-genome bisulfite sequencing to assay genome wide CpG methylation. We combined these results with previously available chromatin immunoprecipitation followed by high-throughput sequencing (ChIP-seq) data for tissue-specific and ubiquitous transcription factors. Specifically, we reanalyzed ChIP-seq data for CTCF, CEBPA, HNF4A, in all five species; FOXA1 in all species but macaque; and ONECUT1 in all species but dog. This allowed us to determine the CpG methylation patterns at transcription factor binding regions, and compare their evolutionary conservation across mammals (Figure 1A). We profiled the methylation of 39-57 millions CpGs in each species at an average of 6-15X coverage (Supplementary Table 1), which accounted for 82-97% of all genomic CpGs (Figure 1B). As previously reported [30],the distribution of genomic CpG methylation is bimodal, with the highest between 80-100% methylation and the lowest between 0-10% (Figure 1C). These coordinated datasets enabled us to investigate relationships between DNA methylation and transcription factor binding.

**Figure 1:**
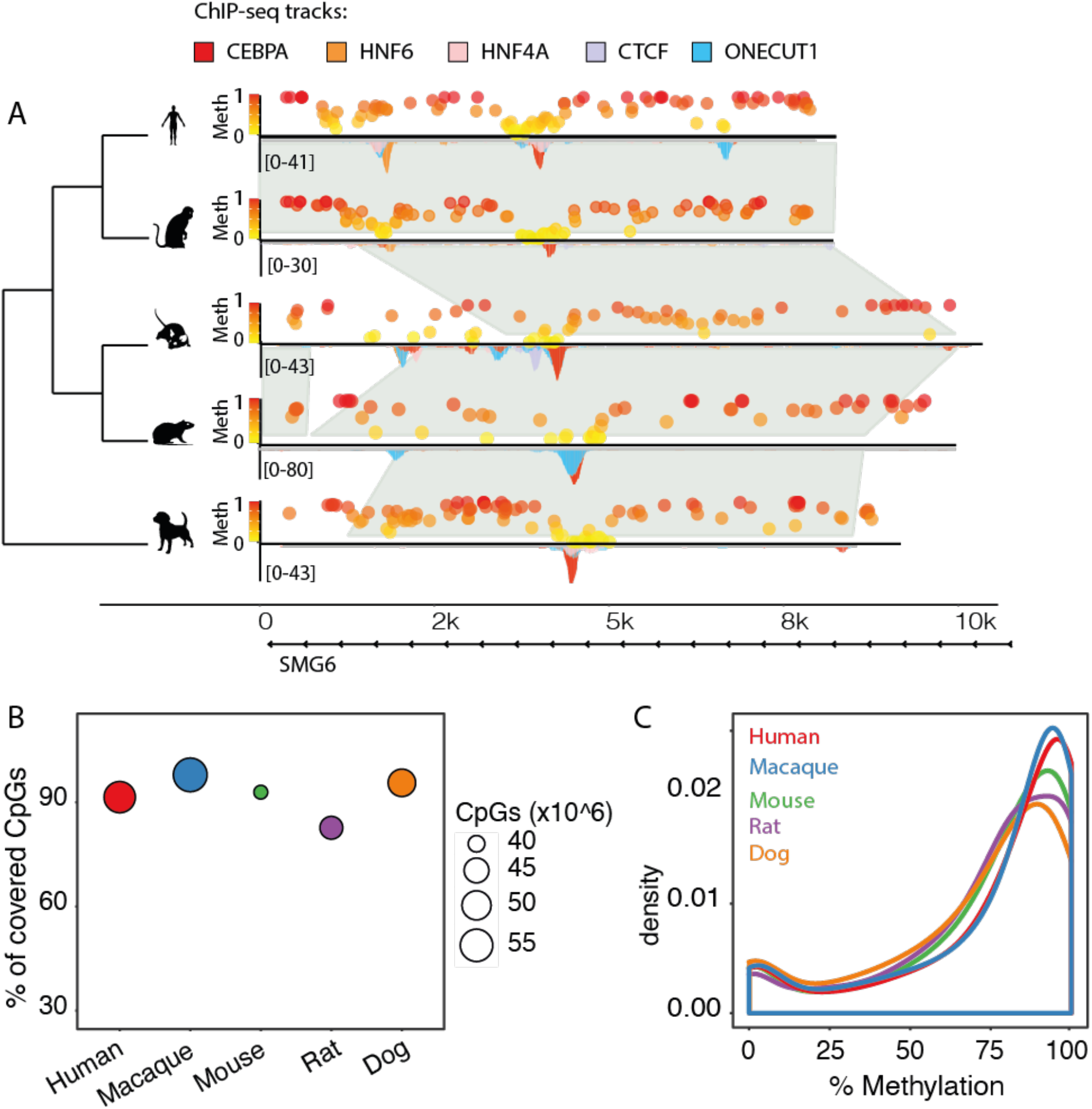
Experimentally mapping methylomes across mammals. **A)** Example cross species whole genome alignment of the SMG6 gene. For each species levels of CpG methylation assayed with bisulfite sequencing are shown above the region, and binding of five transcription factors tracks (CEBPA, CTCF, FOXA1, HNF4A, ONECUT1) assayed through ChIP-sequencing are shown below. **B)** Genomic coverage of WGBS data in each species. The y-axis shows the percentage, while the radius of each point denotes the total number of CpGs covered on the forward and reverse strands. **C)** Genome wide CpG methylation density distributions for each species. All distributions are bimodal, with the vast majority of CpGs hypermethylated.

### Transcription factor binding regions have intermediate CpG methylation

We explored the presence of CpGs and their methylation levels at transcription factor binding regions (TFBRs) and binding sites (TFBSs). Using the ChIP-seq data, we first identified TFBRs by calling peaks with MACS2 [37], and normalized their length to the average peak length estimated separately for each transcription factor and species (Supplementary Table 2, Methods). TFBSs were defined as the DNA sequence where the relevant TF binding motif mapped closest to ChIP-seq peak summit (Methods). We found that around 80% of binding regions harbor at least one CpG in the close proximity of the binding site (Figure 2A). TFBRs frequently have between three and five CpG sites (Figure 2B). However, only a considerably smaller fraction of regions contains a CpG at the binding site (Figure 2A). CTCF is an exception in that approximately 30% of binding sites contain one or more CpGs (Figure 2A). This is also reflected in the canonical motif logos -CTCF has more high-scoring CpGs instances in the position weight matrix (Supplementary Figure S1A). Taken together, transcription factor binding regions commonly contain CpGs, but rarely at the binding site. This suggests that the closely surrounding region has a few key sites that could be methylated and thus potentially affect binding through direct steric hindrance or interference with a binding partner.

**Figure 2:**
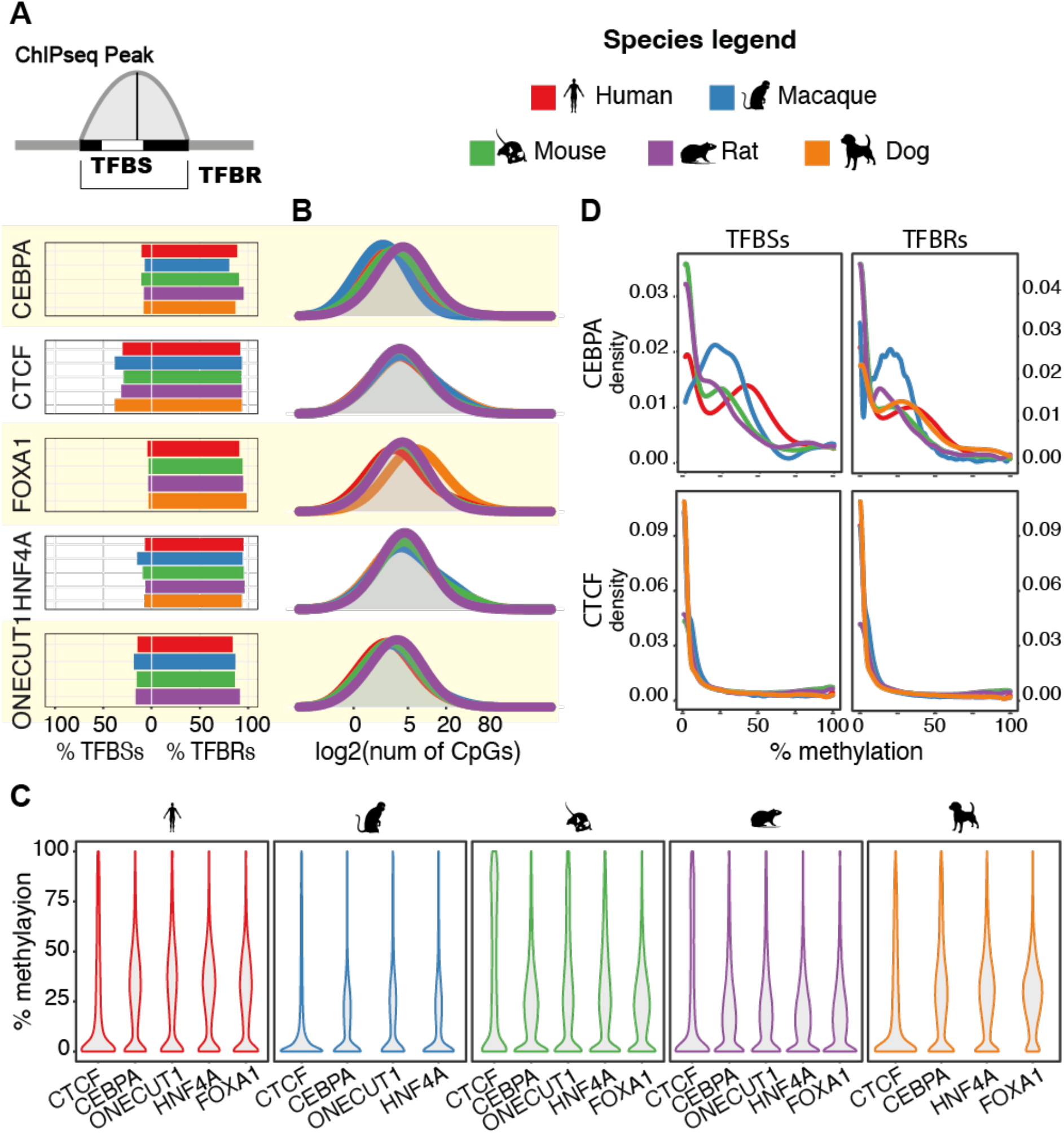
Low and intermediate methylation are signatures of TFBRs. **A)** Percentage of TFBRs and TFBSs harboring at least one CpG for each TF and species. Most TFBRs contain CpGs, but rarely at the TF binding site. **B)** Density distributions of CpGs within TFBRs. TFBRS generally have between three and five CpGs. **C)** Average methylation levels of TFBRs. In all species and most TFs the distributions are bimodal, with the highest mode within intermediate levels of 5mC. CTCF has unimodal distributions across all species. **D)** CpG methylation density distributions at TFBRs and TFBSs. All distributions are bimodal, except for CTCF which has unimodal distributions in all species.

The average methylation across TFBRs is a bimodal distribution (Figure 2C), with both modes below 40%. This differs from the genomic background bimodal distribution (Figure 1C) and suggests that most binding regions are either depleted of methylation or have intermediate methylation levels. Intermediate methylation (IM) is also observed when considering CpG methylation density distributions at binding regions and binding sites (Figure 2D, Supplementary Figure S1B). These distributions are bimodal and confirm that a considerable fraction of CpGs take up intermediate methylation levels, and that IM is not simply an artefact of averaging across the regions. Interestingly, hypermethylation occurs more commonly in CEBPA TFBS than in the wider CEBPA TFBR (Figure 2D). This supports previous observations that CEBPA may bind to methylated motifs [34], while there is no such evidence for the remaining factors. CTCF is again an exception: in all species assayed, it has a unimodal distribution with a large fraction of fully demethylated TFBRs and an absence of intermediate methylation (Figure 2C and 2D). Taken together, hypo- and intermediate methylation are signatures of both TFBRs and TFBSs. The wide range of methylation levels we observed most likely reflects the diverse genomic contexts where TF bind, such as promoters or distal regulatory elements [16].

### Transcription factors bind regions of distinct methylation profiles

To explore the methylation patterns of the genomic neighborhood bound by transcription factors, we extended the TFBRs to 1200 bp and found that CpG frequency is higher than at random genomic regions and increases approaching the peak summit (Figure 3A, Supplementary Figure S2). This is consistent across species and factors, although the width and height of the frequency peaks varies between transcription factors. Inversely to CpG frequency, methylation levels are high at 600 bp away and then sharply fall at the binding summit (Figure 3A, Supplementary Figure S2). This shows that TFBRs are characterized by key CpGs in close proximity of the binding site which are predominantly unmethylated when the protein is bound. A few key features stand out from these profiles. First, CTCF exhibits an oscillatory methylation profile which is likely associated to the strong positional pattern of nucleosomes around CTCF binding sites[38][39]. Second, though CEBPA binding sites have overall low methylation, there is a slight increase in methylation at the binding site. This is consistent with the higher density of hypermethylation in CEBPA TFBSs compared to TFBRs shown in Figure 2D. These average profiles demonstrate the most common methylation patterns around transcription factor binding sites. To study the methylation profiles in more detail, we next asked if the average profiles can be further dissected into distinct patterns of local methylation.

**Figure 3:**
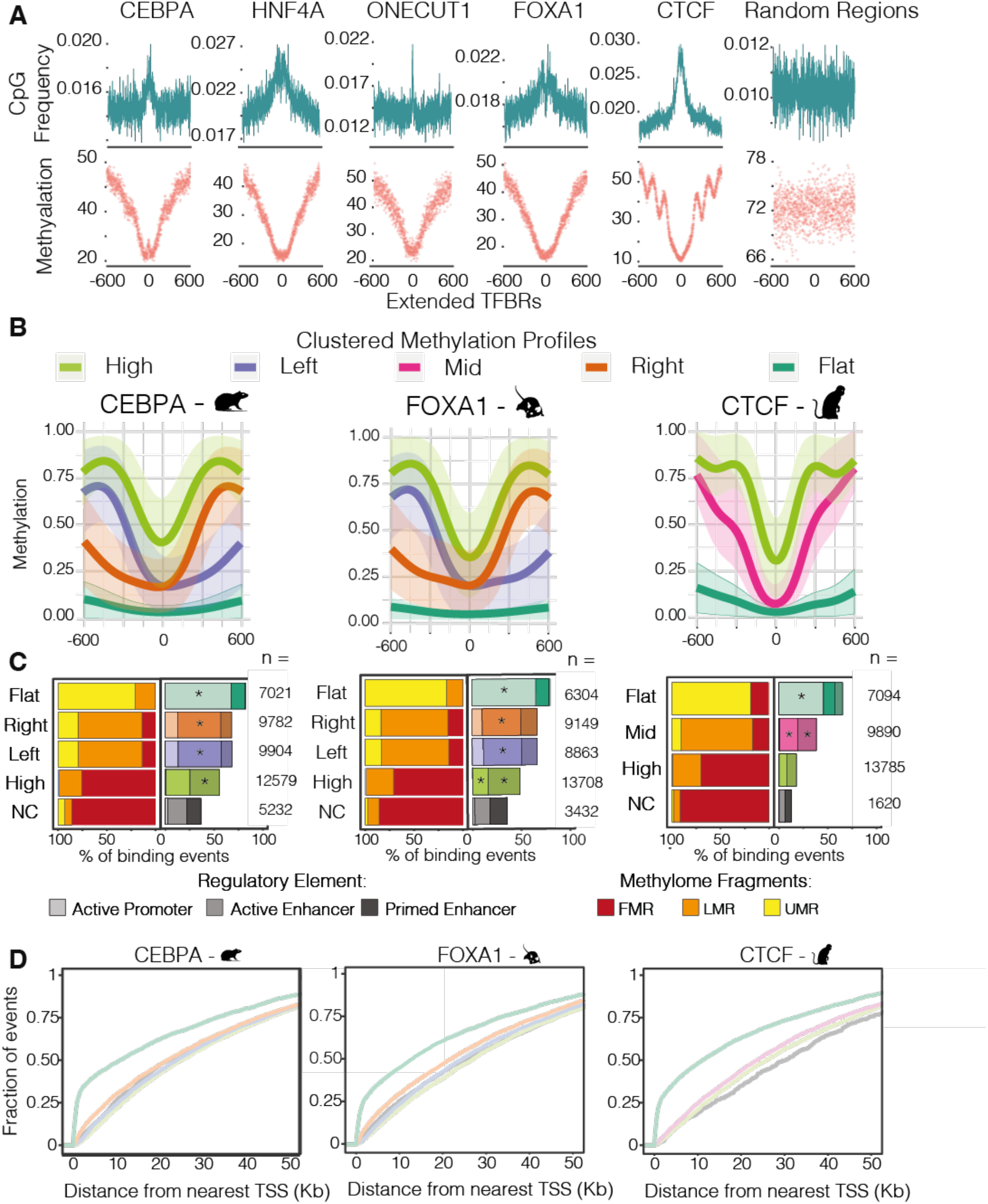
Distinct methylations profiles characterize transcription factor binding regions. **A)** Average 5mC and CpG frequency profiles of human transcription factor binding regions, centered on ChIP-seq peak summits and normalized to 1200bp length. The number of regions classified in each profile is shown in panel B. **B)** Clustered 5mC profiles for rat CEBPA, mouse FOXA1 and macaque CTCF binding regions centered on ChIP-seq peak summits and normalized to 1200bp length. The regions have four types of methylation profiles: “flat” in dark green, “left” and “right” in purple and orange, respectively, (both referred to as “specular” in the text), “high” in light green and “mid”, which is unique to CTCF, in pink. **C)** Annotations of TF binding regions associated to each clustered methylation profiles defined in panel B. On the left, the percentage of TF binding events belonging to each methylation profile located within Unmethylated (UMRs), Lowly-Methylated (LMRs) or Fully-Methylated Regions (FMR) of the genome. On the right, the percentage of TF binding events in each 5mC profile that are annotated as active promoters, active enhancers, or primed enhancers. Asterisks indicate that the annotation category is significantly enriched (z-test with Bonferroni correction, * = p-values << 0.05) **D)** Cumulative distributions of the distance of each TF binding region from the nearest transcription start site, grouped by methylation profiles defined in panel B.

To investigate multiple coexisting DNA methylation patterns, we derived the methylation profiles of extended TFBRs using generalized linear model regression and clustering with the BPRMeth R package [40]. According to the best fit model, all extended TFBRs cluster into three or four prototypical methylation profiles with discernible patterns (Figure 3B and Supplementary Figure S3, Methods). These profiles are very similar across transcription factors and species, and we named them according to their features. The “high” clusters are the most abundant for all factors (Figure 3C and Supplementary Figure S4), they have high methylation 300-500 bases from the binding site and show a narrow drop to intermediate methylation levels at the center (Figure 3B and Supplementary Figure S3). All transcription factors but CTCF were assigned right and left “specular” profiles (so named because they are mirror images of each other) that have 70% methylation at one end of the profile, a drop to 20% methylation at the binding site, and low methylation maintained to the opposite end of the profile (Figure 3B and Supplementary Figure S3). The right and left specular profiles account for about 40% of binding regions (Figure 3C). The last “flat” profile comprises of the smallest group of binding regions (Figure 3C), and is characterized by wide regions of complete demethylation (Figure 3B and Supplementary Figure S3). Only a few thousands binding regions could not be used for clustering due to a small number of CpGs (i.e. less than four), and were hence named non-classified (“NC”). Specular methylation clusters were not observed for CTCF, instead it has an intermediately methylated cluster, the “mid” cluster. The mid is similar to the high cluster, but with a steeper drop in methylation and complete demethylation at the binding site (Figure 3B and Supplementary Figure S3). These prototypical profiles are reproducible across both species and transcriptions factors (Supplementary Figure S3). Taken together, clustering classification provides a robust approach to group transcription factor binding regions according to their distinct and conserved methylation patterns.

### Methylation profiles have distinct contexts and functions

DNA methylation levels and CpG density are associated with different regulatory contexts [41], therefore we next explored if the prototypical methylation profiles associate with distinct regulatory functions. To test the association with regulatory contexts, we annotated TFBRs using available active promoter (marked by histone 3 lysine 4 trimethylation (H3K4me3) and histone 3 lysine 27 acetylation (H3K27ac)), active enhancer (marked by histone 3 lysine 4 monomethylation (H3K4me1) and H3K27ac) and primed enhancer (H3K4me1 only) calls determined by ChIP-seq [42] (Figure 3C right side and Supplementary Figure S4A). To explore their genome-wide methylation context, we annotated TFBRs according to their occurrence in Un-Methylated (UMRs), Lowly-Methylated (LMRs) and Fully-Methylated (FMRs) Regions ([26], Methods) (Figure 3C left side and Supplementary Figure S4B).

A significantly high proportion (around 70%) of TFBRs with the flat profile overlap with active promoters (Figure 3C right side and Supplementary Figure S4B). These are mostly found within UMRs, very close to transcription start sites (TSSs), and 35% overlap an annotated TSS (Figure 3D and Supplementary Figure S5). In fact, many of these regions are also found within CpG islands (Supplementary Figure S6B). This is consistent across all species and for all transcription factors (Supplementary Figure S6B). This shows that TFBRs of the flat clusters are largely promoter regions. The right and left specular profiles comprise of similar numbers of binding regions (Figure 3C), have comparable methylation levels (Supplementary Figure S7A) and similar annotations (Figure 3C and 3D, Supplementary Figure S4 and S5), clearly underlying the same regulatory contexts. Therefore, for further analyses we grouped these two profiles together. Most TFBRs of the specular profiles are significantly found in LMRs and enhancers, with only a small fraction overlapping active promoters. Similarly, the high profile comprises of TFBR significantly overlapping enhancers, but predominantly in fully methylated regions (FMRs). The non-classified (NC) TFBR are found in FMRs and about 40% of them were annotated as enhancers. The enhancer regions of the specular, high and NC groups are equally far from TSSs (Figure 3D and Supplementary Figure S5) and do not overlap with CpG islands (Supplementary Figure S6). Generally, TF binding events associated with the different methylation profiles show similar binding intensities, measured by the ChIP-seq signal fold enrichment distributions (Supplementary Figure 7). However, binding events associated to the high profiles had significantly lower fold enrichment scores than the specular profiles (Supplementary Figure 7). Next, we compared CpG densities associated to the methylation profiles and found that flat profile TFBRs are the richest, while those with the high profile have significantly fewer CpGs (Supplementary Figure S6A), showing that the profiles have unique CpG densities. We show that prototypical DNA methylation profiles distinguish between unmethylated CpG-rich promoter regions, lowly-methylated enhancers with intermediate CpG density, and highly-methylated CpG-poor enhancers.

CTCF has exceptionally few overlaps with regulatory regions, except for TFBRs of the flat profile which have comparable annotations to the other transcription factors. 25% of TFBRs with the mid profile, found only for CTCF, and 10% with the high profile overlap primed or active enhancers. This overlap is too small to confidently assign CTCF TFBRs of these profiles to regulatory contexts. Moreover, TFBRs with a mid profile typically show higher binding intensity (Supplementary Figure 7). Given that CTCF not only has a role in regulation, but also in genome stability and architecture, the low overlap with promoters and enhancers likely reflects methylation landscapes of alternative chromatin contexts.

Taken together, transcription factors bind within different chromatin contexts that are marked by specific DNA methylation profiles which are deeply conserved in mammals.

### DNA methylation levels are coupled to transcription factor binding divergence

We investigated the evolutionary conservation of DNA methylation patterns across species and its association with transcription factor binding divergence. First, we leveraged the EPO multiple species alignments from Ensembl version 98 [43] to define regions orthologous to TFBRs by projecting their coordinates onto the other species’ genome (Figure 4A, Methods). Next, we compared DNA methylation levels across TFBRs and their orthologous counterparts to check for TF binding in the orthologous region. Typically, orthologous regions that do not bind a TF have higher methylation levels than those that do, with average medians of around 75% and 20%, respectively (Figure 4B). The orthologous unbound regions have hypermethylation levels comparable to randomly selected genomic segments (Figure 4B), and to those previously described for the non-regulatory portion of the genome [23]. Next, we leveraged the parsimony principle and the structure of our phylogenetic tree to subset orthologous regions into evolutionary binding losses and gains (Figure 4A, Methods).

**Figure 4:**
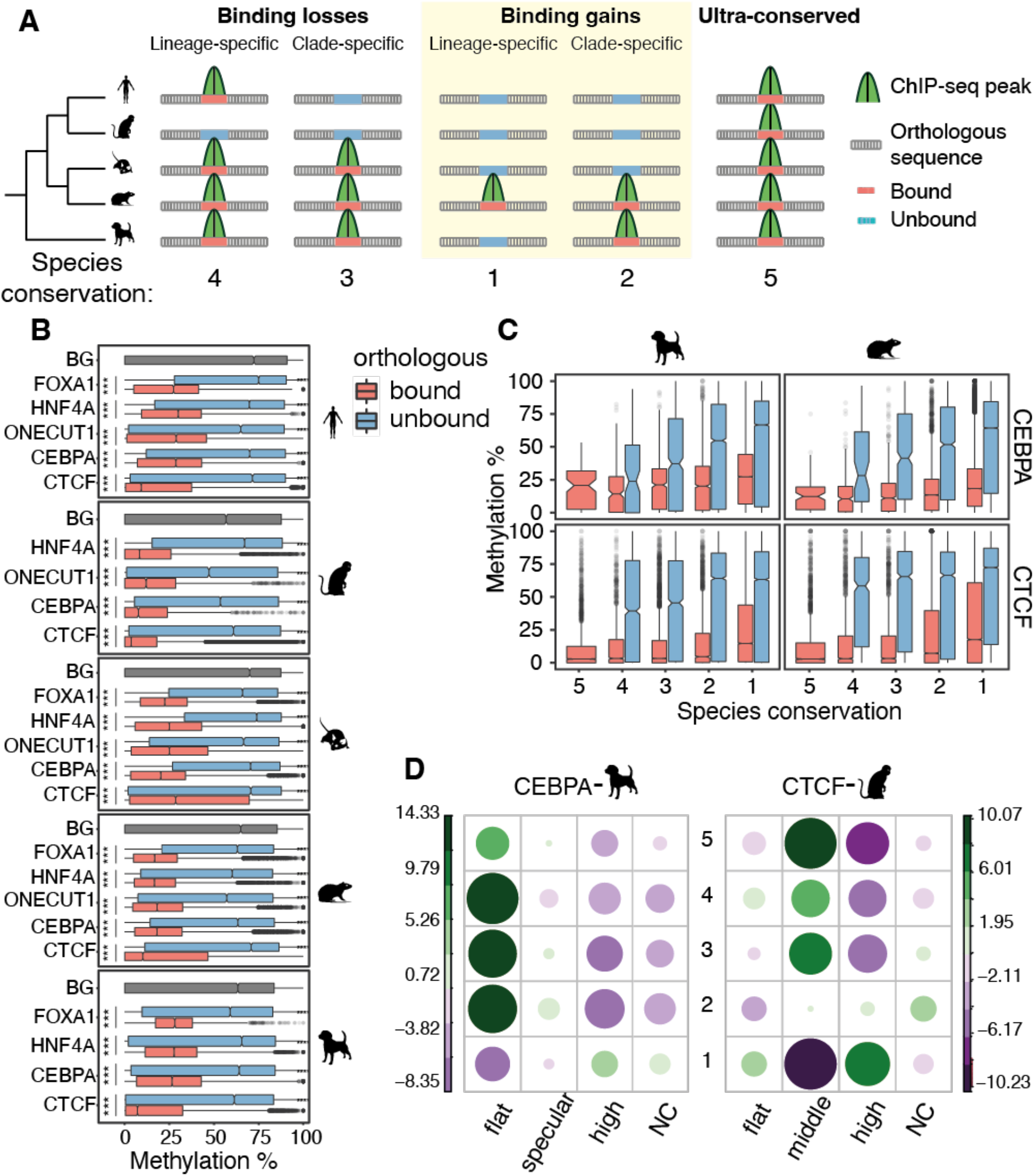
Coevolution of methylation and TF binding in mammals. **A)** Schematic representation of the phylogenetic parsimony approach (adapted from [6]) to define species conservation categories and number of species with binding conservation. Briefly, TF binding events were first aligned and compared across species, then divided using parsimony in lineage- and clade-specific binding losses, and lineage- and clade-specific binding gains. Regions with experimentally determined binding in the species were called as orthologous bound, and those without binding unbound. Ultra-conserved binding events were defined as those bound across all species. Below, examples of corresponding degrees of species conservation defined by the total number of species that share a TF binding event. **B)** Average 5mC level distribution of bound regions, orthologous unbound regions and genomic background (BG), with significant differences marked with asterixis (Wilcoxon test with Bonferroni correction, *** = p-value <= 0.001). **C)** Average 5mC distributions within CEBPA and CTCF bound and orthologous unbound regions divided by species conservation categories defined in panel A (Jonckheere-Terpstra trend test, p-values < 2.2e10^6), shown for dog and macaque. **D)** Relationships between species conservation and 5mC profiles. Balloon plots show Pearson’s residuals from an association analysis between 5mC profiles and TF binding conservation categories for dog’s CEBPA and macaque’s CTCF TF binding events.

We found that the difference between methylation levels of bound and unbound orthologous regions is more pronounced for binding losses than binding gains (Supplementary Figure S8A). These results show coordinated evolution of DNA methylation and TF binding, and suggest that orthologous regions which lost a binding event over evolution are reset back to the hypermethylated levels of the non-regulatory genome.

We further investigated whether there is a correlation between DNA methylation and the degree of TF binding conservation (Figure 4A). Lineage-specific binding events (i.e. those bound in only one of the species studied) have intermediate levels of methylation; as the number of species sharing a binding event at orthologous locations increases, the methylation level decreases (Figure 4C, Supplementary Figure S8B). DNA methylation is inversely correlated to the degree of species conservation, even for orthologous unbound regions. These results show that DNA methylation co-evolves with the binding divergence of specific TFs.

Next, we explored the prevalence of DNA methylation profiles across TFBRs with different degrees of binding conservation. We found a highly significant association between species conservation and profile assignments of TFBR (chi-square test, p value < 0.05). Figure 4D and Supplementary Figure S9 estimate the strength of the association through Pearson’s residuals (Methods), and shows that the high and flat profiles associate most strongly with species conservation, although in opposite directions. Specifically, lineage-specific binding events are negatively associated with the flat profile, and the higher the species conservation the stronger the association. TFBRs with high profiles show the opposite trend; only the lineage-specific binding events have a positive association between species conservation and the profile, while higher degrees of conservation contribute negatively. The specular profiles contribute mildly to species conservation. CTCF TFBRs have different associations: those with flat profiles only mildly contribute to correlation with higher species conservation, instead those with mid profiles contribute more strongly. The association of specific methylation profiles to varying degrees of species conservation can in part explain the co-evolution of TF binding and DNA methylation.

In conclusion, DNA methylation is coupled with TF binding divergence at different levels. Bound regions are more highly methylated than orthologous unbound regions. Furthermore, the methylation of both bound and unbound regions tracks with the degree of species conservation. Finally, different methylation profiles are associated with high and low species conservation, indicating that their regulatory contexts might contribute to evolutionary coupling of TF binding and DNA methylation.

## DISCUSSION

To explore the co-evolution of DNA methylation and TF binding, we combined newly generated bisulfite-sequencing experiments and matched publicly available ChIP-sequencing data for five transcription factors in five mammals. These datasets allowed us to determine the spatial variation of DNA methylation across transcription factor binding regions and characterize the genomic contexts that establish distinct DNA methylation patterns. We leveraged interspecies differences that arose over 96 million years of evolution among the five species and revealed coordinated evolution between transcription factor binding divergence and DNA methylation patterns.

### CpG methylation levels in transcription factor binding regions depend on the genomic context

The extent of DNAm’s role in modulating TF binding through changes of affinity towards their target sequences is still not clear [44] [31]. Our data show that only a small subpopulation of TFBSs contain a CpG and thus could be directly affected by 5mC. Considering the instability of methylated cytosines [45], this suggests that CpGs may be generally negatively selected at TFBSs and those present could be protected from mutagenic processes through other mechanisms, but further investigation is necessary to confirm this hypothesis.

The wider genomic context surrounding the investigated TFBRs more often harbor CpGs than their TFBSs. Their methylation state is less likely to directly disrupt the TF binding site, however it can still affect binding through processes such as steric hindrance or the recruitment of chromatin remodelers. Our results therefore suggest that local demethylation in TFBRs is rarely due to the direct competition between transcription factors and DNAm levels [46],[16]. Furthermore, the intermediate methylation and complete demethylation that we observed at TFBRs are consistent with a recently published model describing distinct methylation dynamics between different genomic contexts [25]. For example, it showed that intermediate methylation at distal regulatory regions is the result of an increased rate of passive demethylation and variable rates of *de novo* methylation.

To enhance interpretability across genomic contexts, we further described local methylation patterns within TFBRs and their genomic surroundings using generalized linear model regression and clustering [28] [40]. We revealed that methylation patterns of TF binding regions can be summarized in three prototypical profiles, and reflect their genetic and chromatin contexts. The profile with low levels of methylation throughout (i.e. flat) was typical of CpG-rich promoter regions, and is likely the results of H3K4me3’s inhibition of *de novo* methylation [47]. On the other hand, profiles with intermediate levels overall were enriched within distal regulatory elements marked by H3K4me1. The high profiles had intermediate to high methylation and mostly occurred in CpG-poor enhancers. The specular profiles had low to intermediate methylation and were also marked by the active histone mark H3K27ac. Thus, different types of regulatory regions can be discriminated solely based on 5mC patterns.

Taken together, our results suggest that local methylation levels are determined through competition among a wider number of context-dependent regulatory players such as transcription factors, chromatin remodelers and DNA methylation effectors.

### Coevolution of 5mC and transcription factor binding

Although most transcription factors bind extremely conserved DNA motifs in mammals, their genome-wide binding patterns are highly divergent between species [3][6]. Our study reveals that DNAm follows inter-species divergence of cis-regulatory activity. Specifically, 5mC levels are low at TF bound regions, but they increase at orthologous locations after binding loss to levels of non-regulatory intergenic CpGs. Given that DNA methylation broadly mimics occupancy of various TFs [8] [48], the detected gains of methylation may be indicative of complete regulatory turnover of the orthologous region.

Within each genome, we showed that 5mC levels are inversely proportional to the number of species with conserved binding. This is true for all orthologous regions regardless of whether they are bound, though unbound regions have higher methylation values on average. These results can partly be explained by enhancers evolving more rapidly than promoters [49],[50]: binding sites with flat methylation profiles characterize promoters and have low within-species methylation, while those with high methylation profiles characterize enhancers and have higher within-species methylation.

We show that the genomic context partially explains the evolutionary relationship between 5mC and TF binding divergence, but more elaborate models are needed to define the rate of DNAm turnover within these contexts.

### CTCF has distinct CpG methylation profiles and evolution

Our analyses demonstrate that CTCF profoundly differs from the other TFs investigated in its DNA methylation landscape and evolution. Unlike other TFs, 30% of CTCF binding sites contain at least one CpG and CpGs close to the ChIP-seq peak summit are overall more stably and lowly methylated. At the same time, CTCF binding events are found in hypermethylated contexts more often as 5mC rapidly increases around the binding site. Aside from CTCF TFBRs found within CpG-rich promoters (i.e. flat profiles), the remaining CTCF TFBRs have different profiles that are not linked to promoter or enhancer elements. These differences in 5mC profiles may be explained by their different functions: CEBPA, HNF4A, ONECUT1 and FOXA1 are transcription factors involved in the activation of key liver genes [35], while CTCF is a multifunctional and ubiquitous transcription factor involved in gene activation but also in determining three-dimensional genome structure, insulation, and gene repression [36,51].

Finally, CTCF’s 5mC profiles evolve differently than those of the other TFs. Specifically, the flat CTCF profile, despite being enriched with promoters, is not associated with high species conservation. This may be due to the previously described redundancy of CTCF near functionally important sites (Kentepozidou et al. 2020), which may buffer the loss of a CTCF binding event in one species through turnover. This suggests that CTCF’s binding events within the flat cluster are under less stringent evolutionary pressure than the wider promoter region. On the other hand, the CTCF-specific methylation profile (i.e. mid) is strongly associated with high species conservation, depleted at lineage-specific binding events, and may be subjected to high evolutionary pressure. This points to an important role for these binding sites, but further work is necessary to characterize their function.

## METHODS

### Publicly available data

All ChIP-seq data are publicly available and were retrieved from ArrayExpress (https://www.ebi.ac.uk/arrayexpress). CTCF ChIP-seq data for all species can be downloaded under accession number E-MTAB-437. HNF4A, ONECUT1, FOXA1 and CEBPA ChIP-seq data for all species can be retrieved under accession number E-MTAB-1509. We used all the available experiments except ONECUT1 from dog, due to the lack of replicates. ChIP-seq of histone modifications and processed regulatory region calls can be accessed in ArrayExpress with accession number E-MTAB-7127.

### Tissue preparation

Mammalian liver samples were extracted post-mortem, perfused with PBS, and flash-frozen in liquid nitrogen. Tissues were prepared immediately post-mortem (typically within an hour) to maximize experimental quality, and were kept on ice until processed to minimize potential DNA degradation. Total genomic DNA was extracted from each sample with commercial reagents and following manufacturer guidelines (Qiagen, DNAEasy Blood&Tissue kit). Details on origin, number of replicates, sex and age for each species’ sample are in Supplementary Table 1.

DNA from at least two independent biological replicates from different animals was prepared for each species. Wherever possible, livers from young adult males were used. Samples of healthy liver tissue from humans were obtained from the Addenbrooke’s Hospital at the University of Cambridge under license number 08-H0308-117 ‘‘Liver specific transcriptional regulation’’. Mouse samples were obtained from the Cambridge Institute under Home Office license PPL 80/2197.

### Whole-genome bisulfite sequencing (WGBS) protocol

Mammalian DNA was subjected to bisulfite conversion using Epimark CT conversion kit using Agilent and/or Epimark polymerase (Supplementary Table 1). Subsequently libraries were prepared using NEBNext Ultra DNA library preparation kit and sequenced using an Illumina massively parallel sequencer.

## Genome resources

All genomes were downloaded from the Ensembl ftp version 98 [52] as toplevel assembly files. We then filtered out patches and scaffolds and retained only assembled chromosomes. The species genome versions used are the following: GRCh38.p13 for human, GRCm38.p6 for mouse, Mmul_10 for macaque, Rnor_6.0 for rat and CanFam3.1 for dog.

### WGBS data processing

Paired-end FASTQ files were trimmed and adapters removed using *TrimGalore!* version 0.6.4_dev [53] with default parameters. We then processed the data using *Bismark* version 0.22.3 [54]. First, we performed in-silico bisulfite conversion of the reference genome, i.e. C→T and G→A conversions, using the *bismark_genome_preparation* script. Next, reads were mapped to each species’ genome by running *bismark* with default parameters. Duplicate reads were removed from bam files with *bismark_deduplicator*, before extracting methylation calls using *bismark_methylation_extractor* with the following parameters: *bismark_methylation_extractor --comprehensive --merge_non_CpG --bedGraph --no_overlap --ignore_r2*. Finally, we generated a coverage file using the script *coverage2cytosine* with the following parameters: *coverage2cytosine --merge_CpG --zero_based*. Methylation calls were considered in downstream analyses only if supported by methylation evidence from at least four CpGs (i.e. minimum four read coverage).

### ChIP-seq data processing

Paired-end FASTQ files were trimmed and the sequencing adapters removed using *TrimGalore!* [53] with default parameters. Trimmed reads were then mapped to each species’ genomes using *bowtie2* version 2.3.5.1 [55] with default parameters. We next called peaks using *MACS2* version 2.1.4 [37] using the narrow peak mode and the *-f BAMPE* parameter. FOXA1 experiments from macaque were removed from further analyses due to a smaller number of peaks called compared to the other species. To call reproducible peaks we found overlap between replicates’ peaks with *bedtools intersect* v2.29.2 [56] and kept those that overlap with at least one base between both replicates. For further analyses, we represented the reproducible peak as the original replicate peak with the strongest signal, as defined by the MACS2 p-value.

### Defining transcription factor binding regions (TFBRs) and their methylation coverage

To define transcription factor binding regions (TFBRs) we normalized reproducible peak sets for length. Specifically, we extended the reproducible peaks from the peak summit equally in both directions until we reached the average total peak length within that species and factor. To calculate the number of CpGs that overlap with TFBRs, we used *bedtools intersect* and *bedtools groupy* to intersect the TFBRs with the methylation coverage file. We repeated the same process to calculate the average methylation level associated with TFBRs, but only considered CpGs covered at least four times.

### Motif discovery and transcription factor binding site (TFBS) annotation

Motif discovery was conducted with the *MEME suite* version 5.0.5 [57]. From each peak set we selected the 500 strongest peaks, i.e. with lowest MACS2 p-values, and restricted them to 100bp centered on the peak summit. From this representative set, we performed de-novo motif discovery for the most significant motif using MEME with the following parameters *-mod oops -dna -revcomp - nmotifs 1*. Next, we identified motif matches to these newly generated motif position weight matrices (PWMs) in each TFBR set with *FIMO* using a p-value threshold *-thresh 0.005* and the option *-max-stored-scores 1000000000*. To define transcription factor binding sites (TFBSs), we kept the motif closets to the peak summit. We calculated the number of CpGs and average methylation levels within TFBSs with the same procedure as for TFBRs.

### 5mC and CpG density profiles

We modelled 5mC profiles with an average methylation approach and using the probabilistic model implemented in the *BPRMeth R* package [40]. We first extended TFBRs to 2Kb centered on the ChIP-seq summit, then intersected these extended regions with methylation coverage files.

To calculate average 5mC profiles (Figure 3A and Supplementary Figure S2), we first created a matrix in *R* version 4.0.1 [58] where each row is an extended TFBRs and the positions denote presence and methylation levels of CpGs covered at least 4 times. For each column we calculated the average methylation level and plotted the results using *ggplot2 jitters* version 3.3.4 [59]. To calculate CpG frequency, we repeated the same process but used all CpGs, regardless of coverage.

To model and cluster 5mC profiles with a probabilistic approach (Figure 3B and Supplementary Figure S3), we inferred profiles using the mean-field variational inference (Variational Bayes) method from the *BPRMeth* R package v1.8.2 [40]. For each species and each transcription factor, we independently optimized the number of radial basis functions (RBFs) – which determine the spatial resolution of the methylation profiles – and the number of clusters. To do so, we used the Bayesian Information Criterion (BIC) and set the number of clusters to three for CTCF and to four for the remaining transcription factors. The combination of parameters selected for each species and transcription factor is shown in Supplementary Figure S3.

### Functional categories of TF binding regions

We further annotated TFBRs into functional categories using regulatory region calls from [42], CpG islands (CGIs) and methylome segments annotations. CpG islands were calculated for each species’ genome with the *EMBOSS cpgplot* v6.6 [60] using default parameters. To segment the methylome in UnMethylated Regions (UMRs), Lowly-Methylated Regions (LMRs), and Fully-Methylated Regions (FMRs) we used the *MethylSeekR R* package version 1.22 [26], setting the FDR cutoff to 5 and the *m* parameter to 0.5. To make the assignments, TFBRs were overlapped with each functional category above using *bedtools intersect* v2.26 (1bp overlap required). In Figure 3C (left side, Supplementary Figure 4B) we show the distribution of TFBRs’ clustered profile assignments between these functional categories. Finally, we defined transcription start sites (TSSs) as the start of every annotated transcripts in GTF files downloaded from the Ensembl version 98 ftp [52]. We used *ggplot2* to plot cumulative distributions of the distance between TFBRs and their the closest TSS (Figure 3D and Supplementary Figure S5).

### Evolutionary conservation of TF binding regions

To make evolutionary conservation calls for TFBRs we used their overlap in whole genome alignments. Specifically, we used the EPO eutherian mammal alignment from Ensembl version 98 [43] to align each TFBR with every other species’ genome and extract the genomic coordinates of these orthologous sequences. Next, we overlapped the orthologous coordinates with transcription factor binding locations of the corresponding TF from the species projected to, and if overlap was found we called the binding conserved. For example, CEBPA binding locations from mouse were first aligned to the rat genome, then the aligned orthologous locations on the rat genome were intersected with CEBPA binding locations from rat. If the projected sequence and the rat CEBPA binding region overlapped with at least 1bp, these were considered conserved between mouse and rat. For each binding event within each species, we then summarized the number of species the binding sequence was alignable to and the number of species the binding sequence was both alignable and conserved.

We categorized binding events in two ways (Figure 4A), the first according to the number of species with conserved binding and the second according to phylogeny. The number of species with conservation was defined irrespective of phylogeny; for example, if the binding event was shared at orthologous locations in exactly three species, that was called a 3-way conserved binding event. The second categorization was based on the phylogenetic relationships between the species. Specifically, we considered binding only shared by mouse and rat exclusive to the rodent clade and binding only in human and macaque exclusive to the primate clade. We further built on the phylogenetic approach using the parsimony method defined in [6] to call binding events as ultra-conserved (if the binding event is shared by all species studied), lineage-specific binding loss (if the binding event is present in all species except one), lineage-specific binding gain (if the binding event is present only in one species), clade-specific binding gain (if the binding event is present in only one clade), and clade-specific binding loss (if the binding event is present in all other species except one clade).

To explore the effect of methylation levels on binding conservation, we intersected the orthologous sequences on each species’ genome with the corresponding methylation coverage files to obtain the number of CpGs and their methylation levels, regardless of whether the corresponding TF was bound in that species. In Figures 4B, 4C and Supplementary Figure S8A, we calculated average 5mC levels for the bound and unbound sequences within each evolutionary category. For example, within a rodent-specific binding gain category (also defined as a 2-way binding event), the bound regions correspond to the orthologous regions where a TFBR was identified in mouse and rat, while the unbound regions correspond to the orthologous locations in the remaining species not bound by the TF.

### Association between 5mC profiles and evolutionary conservation

To test for independence between evolutionary categories of binding conservation and clustered methylation profiles, we first used them to create a contingency table in *R* and then performed a chi squared test using the *chisq.test* function. We extracted the chi squared residuals and plotted the correlation as balloon plots with the *corrplot* R function (Figure 4D and Supplementary Figure S9).

## DECLARATIONS

### Ethics approval and consent to participate

Use of animal tissues in this study was covered by the Animal Welfare and Ethics Review Board, under reference number NRWF-DO-02v2, following the Cancer Research UK Cambridge Institute guidelines.

### Competing interests

P.F. is a member of the Scientific Advisory Boards of Fabric Genomics, Inc. and Eagle Genomics, Ltd.

All other authors declare no competing interests.

### Funding

Funding for this study was provided by Wellcome (WT108749/Z/15/Z, WT202878/Z/16/Z, WT202878/B/16/Z), European Research Council (grants 615584, 788937), and the European Molecular Biology Laboratory.

### Authors’ contributions

M.RI., P.F. and M.RO. conceived and designed the study; M.RI. led and conducted the data analysis; N.W. performed experiments; J.Z. handled and processed data; D.V. and D.T.O. provided tissues; M.RI., P.F. and M.RO. wrote the manuscript; J.T. oversaw the experiments; P.F. and M.RO. oversaw the analyses. All authors read and approved the final manuscript.

## Acknowledgments

We thank Chantriolnt-Andreas Kapourani for helpful discussions about BPRMeth R package usage; Paul Ginno for careful reading of the manuscript; Rachel D. Edgar, Elissavet Kentepozidou, Vasavi Sundaram, Dhoyazan Azazi and Maëlle Daunesse for the support and helpful discussions.

## Additional information

**Supplementary File 1**: all supplementary figures related to this manuscript.

**Supplementary Table 1**: description of the WGBS experiments performed in this study, including the unique identifiers for each library and tissue sample collection details for each species.

**Supplementary Table 2**: table containing the normalized peak lengths for each transcription factor and species.

**Additional Data 1**: Additional details on CEBPA and ONECUT1 binding events. A comprehensive table including transcription factor binding coordinates, genomic and 5mC profiles annotations and their cross-species alignability and conservation

**Additional Data 2**: Additional details on CTCF and FOXA1 binding events. A comprehensive table including transcription factor binding coordinates, genomic and 5mC profiles annotations and their cross-species alignability and conservation.

**Additional Data 3**: Additional details on HNF4A binding events. A comprehensive table including transcription factor binding coordinates, genomic and 5mC profiles annotations and their cross-species alignability and conservation.

---

**Figure S1:**
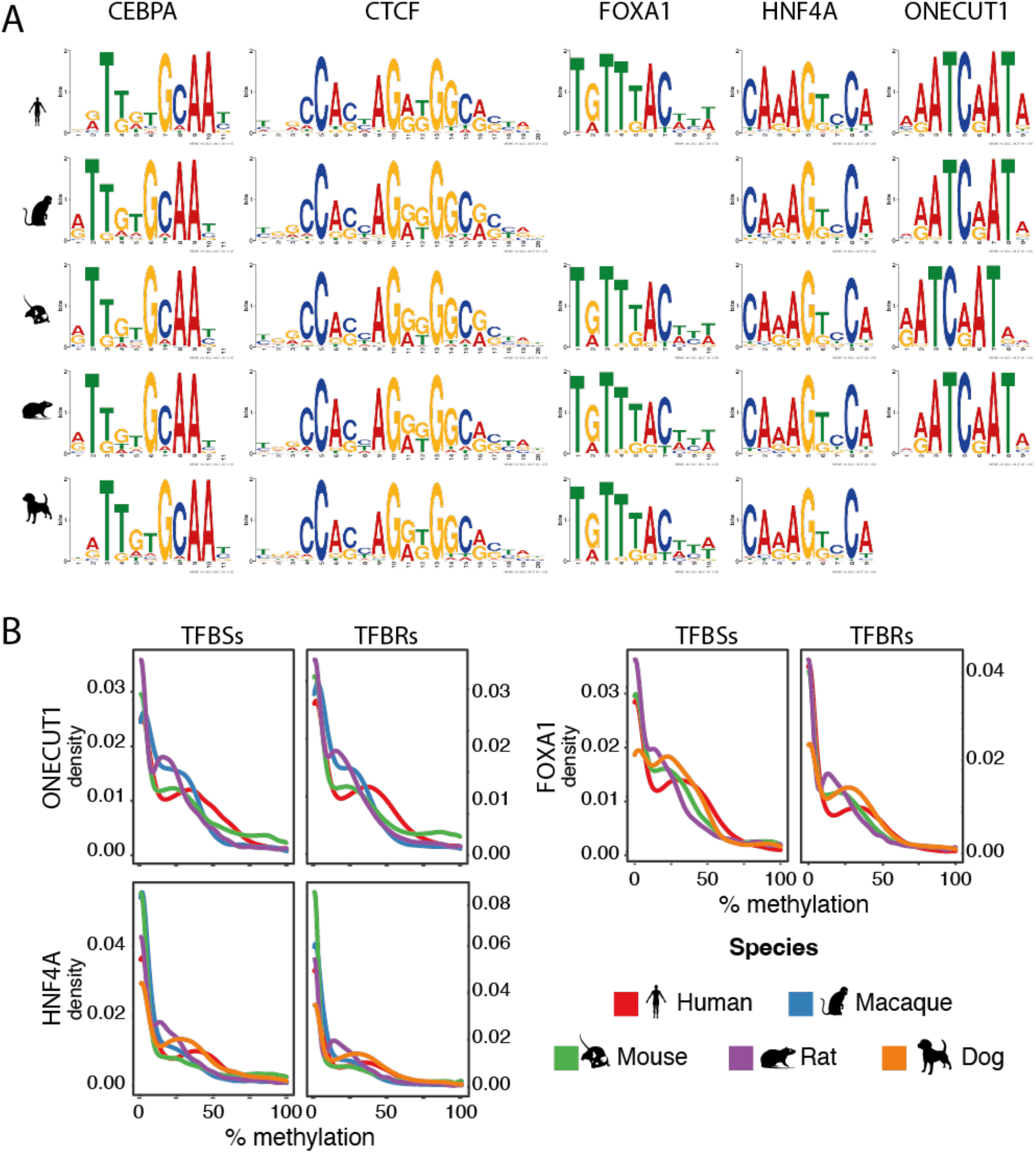
**A)** Position weight matrices (PWMs) calculated de novo for all transcription factors in this study and in all species. **B)** CpG methylation density distributions at TFBRs and TFBSs for ONECUT1, HNF4A and FOXA1 in all species.

**Figure S2:**
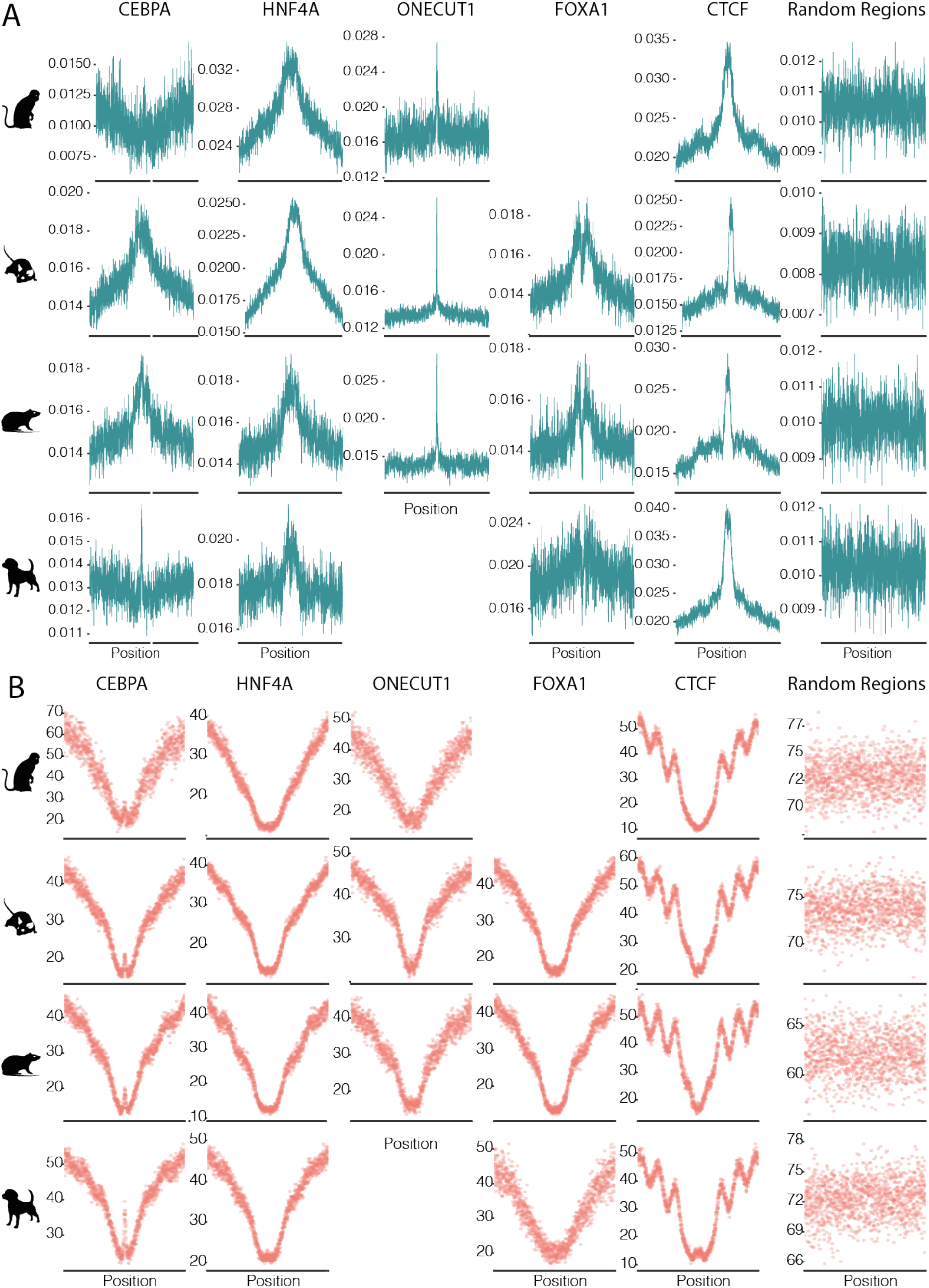
Average CpG frequency (panel A) and 5mC profiles (panel B) at 1200bp wide transcription factor binding regions, centered on ChIP-seq peak summits for all species.

**Figure S3:**
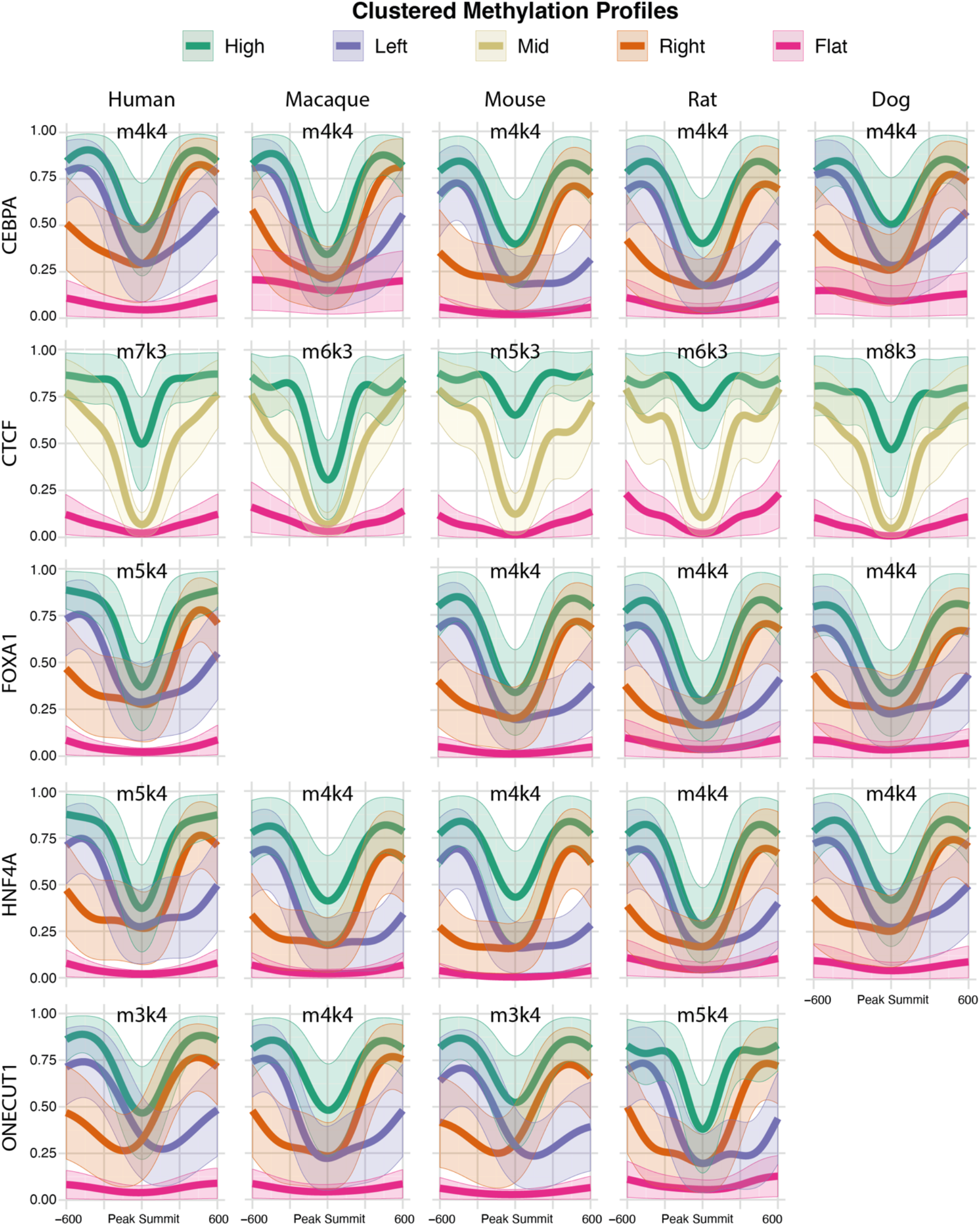
Clustered 5mC profiles for all species and TFs. Above each plot, the optimized M and K parameters are reported.

**Figure S4:**
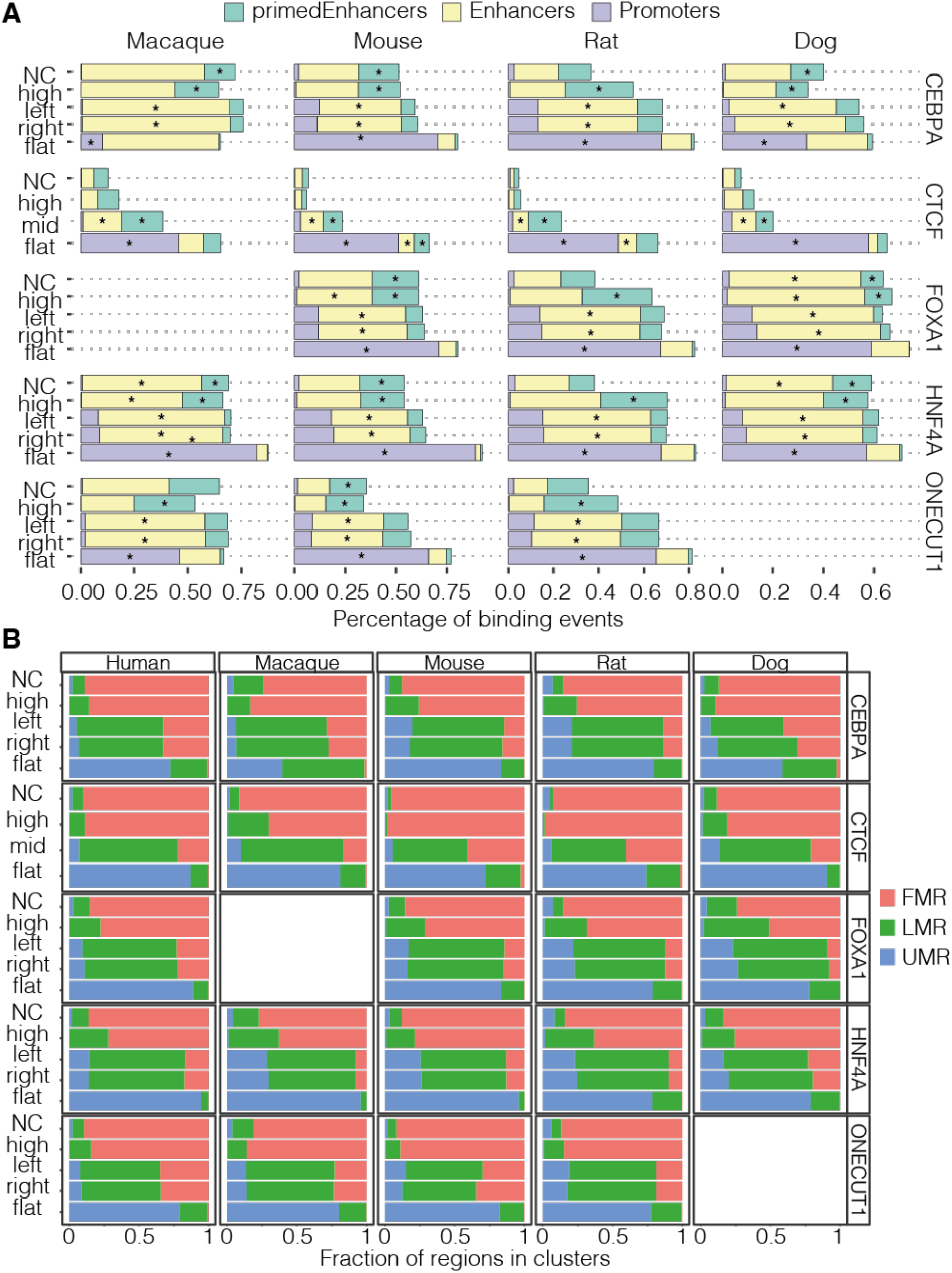
Annotation of TFBRs associated to the clustered 5mC profiles for all species and TFs. **A)** Proportion of TFBRs that overlap with annotated promoters, active enhancers, and primed enhancers. Asterisks indicate enrichment of annotation type (z-test, p-value < 0.05). **B)** Proportion of TFBRs that overlap with annotated UMRs, LMRs or FMRs. FOXA1 ChIP-seq was not available for macaque, and ONECUT1 was not available for dog.

**Figure S5:**
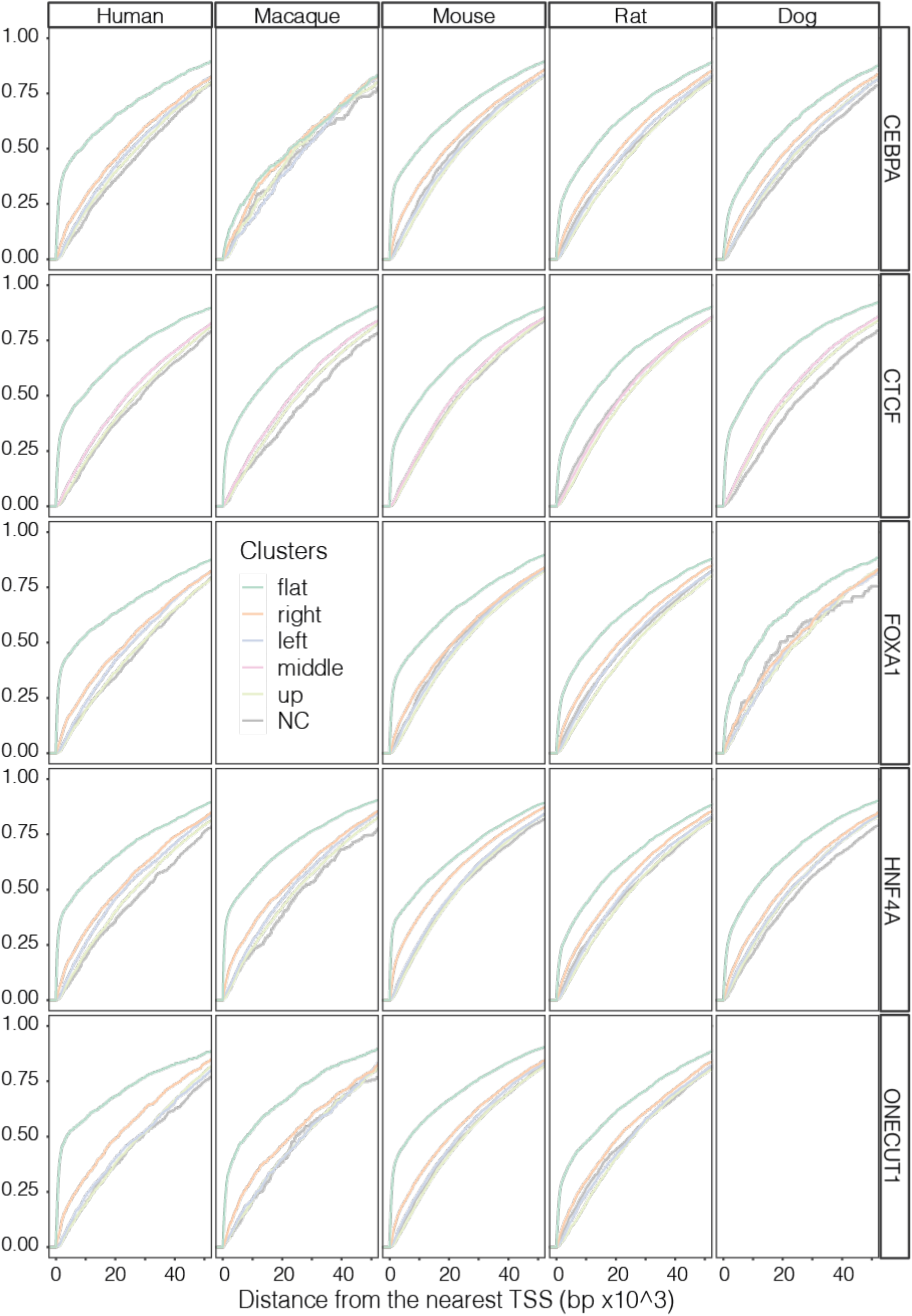
Cumulative distributions of the distance of each TF binding region from the nearest transcription start site, grouped by 5mC profile. FOXA1 ChIP-seq was not available for macaque, and ONECUT1 was not available for dog.

**Figure S6:**
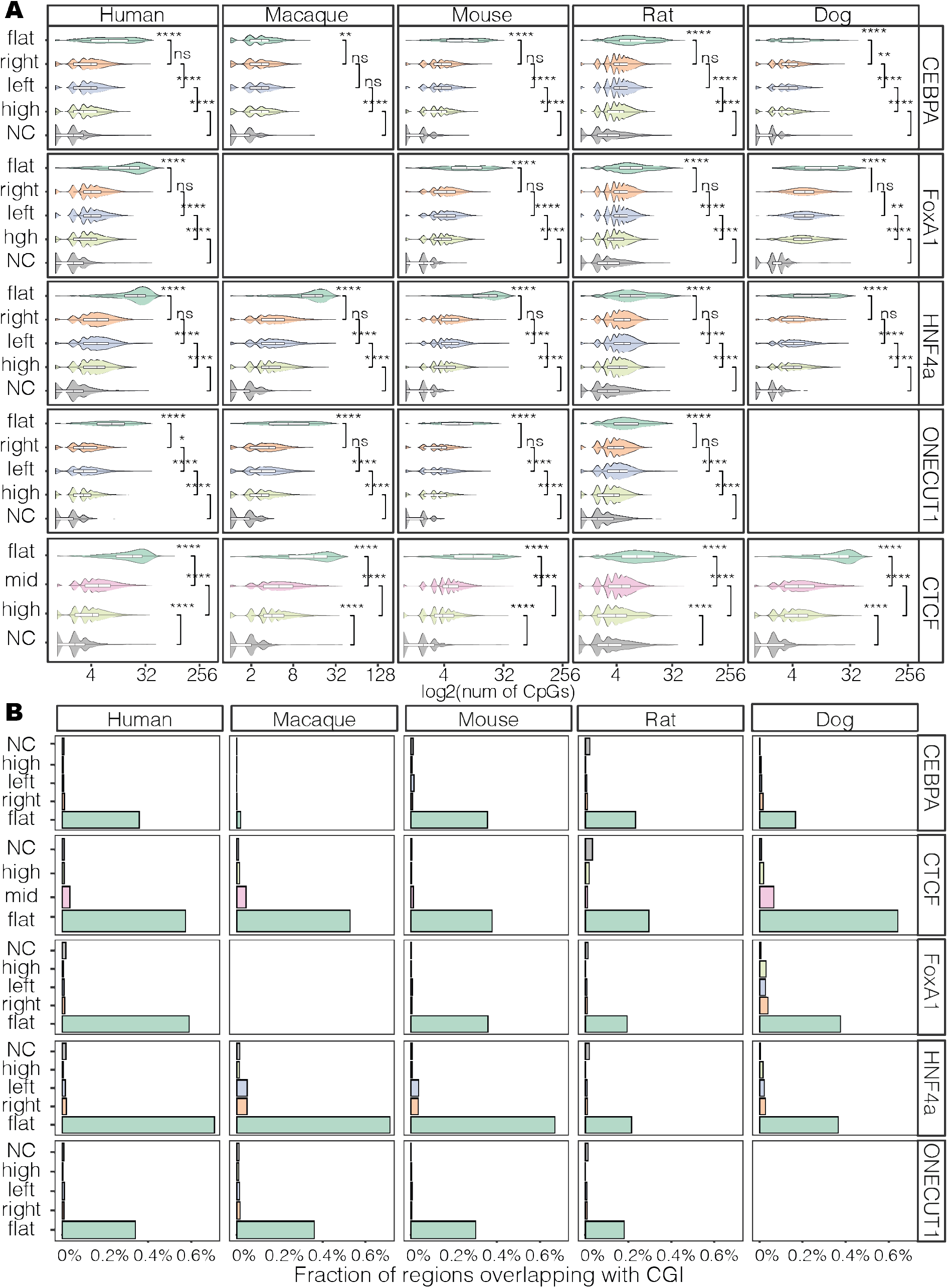
**A)** CpG counts distributions at TFBRs associated with different clustered methylation profiles (Wilcoxon Rank test, p-value < 0.05). **B)** Fraction of TFBRs associated with clustered methylation profiles that overlap with CpG islands (CGI). FOXA1 ChIP-seq was not available for macaque, and ONECUT1 was not available for dog.

**Figure S7:**
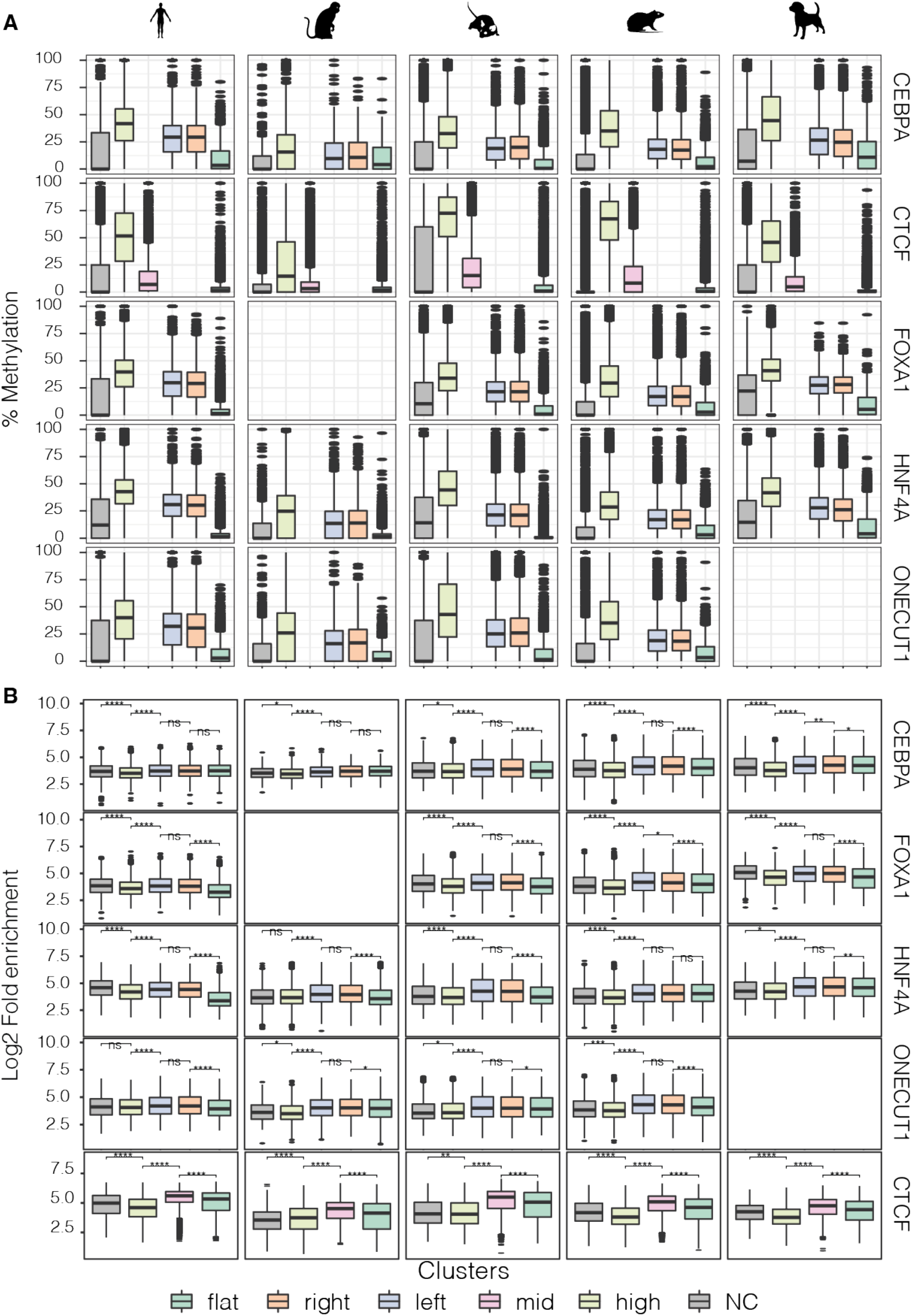
**A)** Average methylation levels distribution of TF binding regions associated with different clustered 5mC profiles. **B)** distribution of fold enrichment values of TF binding regions (ChIP-seq peaks) associated with different clustered methylation profiles (Wilcoxon rank test, ****: p<0.0001, ***: p<0.001, **: p<0.01, *: p<=0.05, ns: p>0.05). FOXA1 ChIP-seq was not available for macaque, and ONECUT1 was not available for dog.

**Figure S8:**
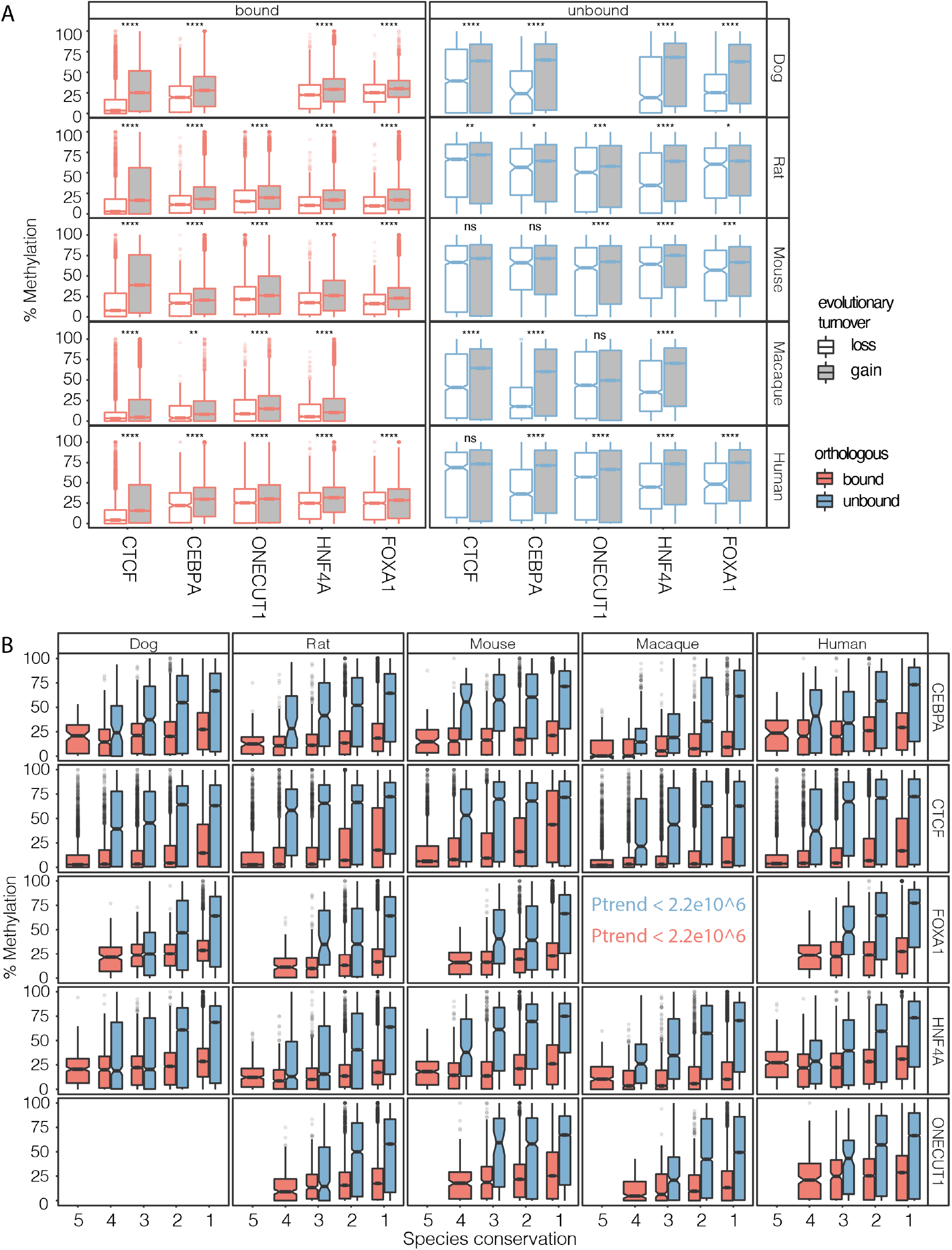
**A)** Average 5mC levels distribution of binding gains sequences compared to binding loss (Wilcoxon rank test, p-value < 0.05; Bonferroni correction). **B)** Average DNA methylation levels distribution of TF bound and unbound regions divided by species conservation categories (Jonckheere-Terpstra trend test, p-values < 2.2e10^6). FOXA1 ChIP-seq was not available for macaque, and ONECUT1 was not available for dog.

**Figure S9:**
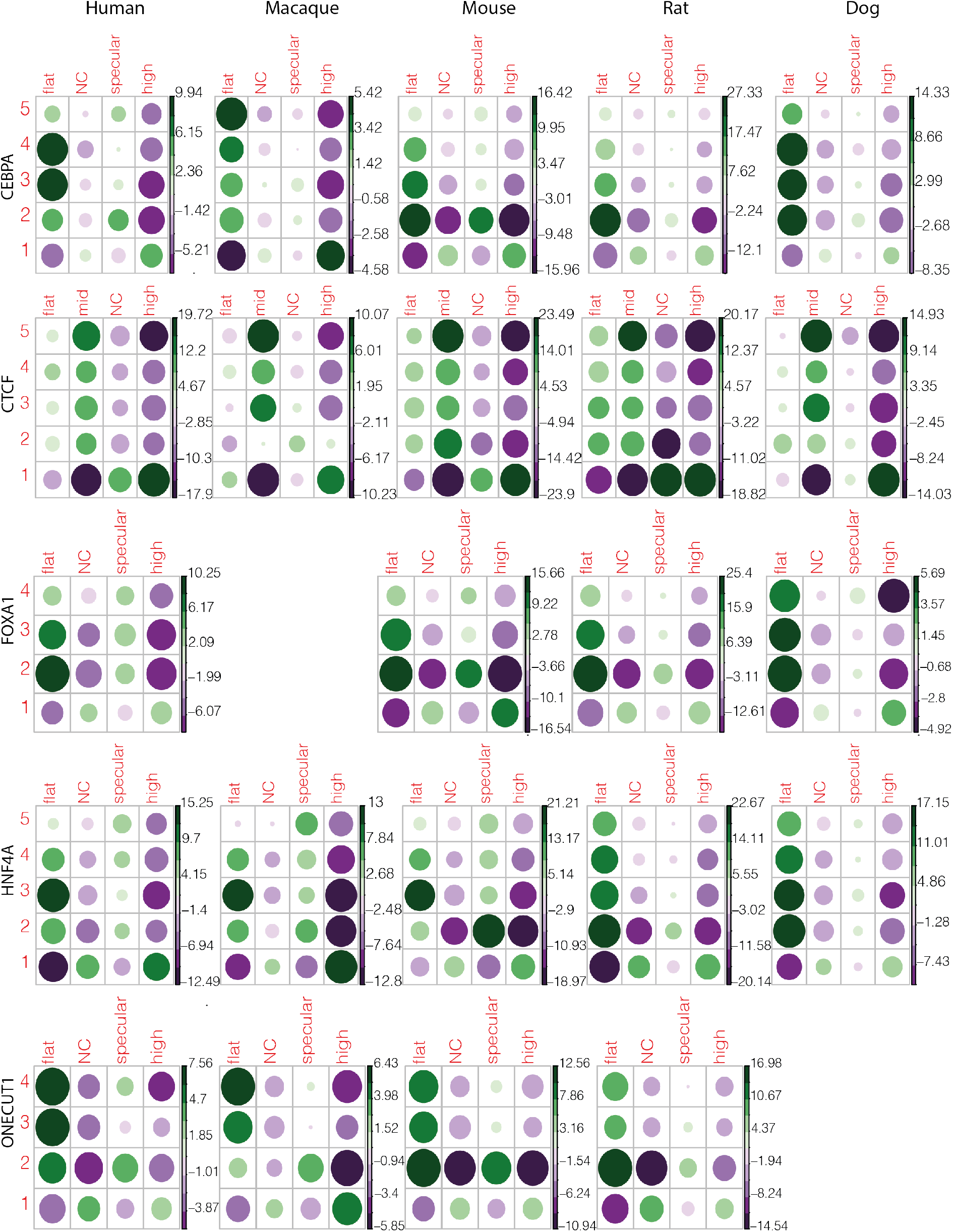
Balloon plots showing Pearson’s residuals from the association analysis between 5mC profiles and TF binding conservation categories for all species and TFs. FOXA1 ChIP-seq was not available for macaque, and ONECUT1 was not available for dog.

