## Supplementary Table 1 for "DNA methylation patterns of transcription factor binding regions characterize their functional and evolutionary contexts": README

README FILE
COLUMN DESCRIPTIONS
"chr": chromosome number
"start": TFBR start coordinate
"end": TFBR end coordinate
"summit": peak summit coordinate
"foldE": peak fold Enrichment (MACS2 output)
"pval": peak p-value (MACS2 output)
"nCpGtot": total number of CpGs within TFBR
"nCpGfilt": filtered number of CpGs (i.e. number of CpGs that have >4X coverage)
"avgMethtot": average methylation across the TFBR, considering all CpGs within the area
"avgMethfilt": average methylation across the TFBR, considering CpGs with >4X coverage
"align": list of species the TFBR aligns to
"num_align": number of species the TFBR aligns to
"bindingCons": list of species with conserved binding at alignable position (if the binding is not conserved, the only species in this list is the species where the binding event was originally assayed)
"num_bindingCons": number of species the binding event is shared with
"cat": conservation category according to parsimony, more general
"label": conservation category according to parsimony, more specific (for example: instead of SpSpecificGain (=cat), here you would have DogSpecificGain)
"binding": either "bound" or "unbound". if "bound", the row is referring to an actual binding event. if "unbound", the row is referring to a projected binding event with no conserved binding (hence, the MACS2 outputs like foldE, summit and pval should be blanlk)
"TF": Transcription factor (ONECUT1 is reported as HNF6)
"Species": Species genome the row refers to
"SpeciesFrom": if binding=bound, this is the same as Species. if binding=unbound, this field tells you which species the binding event was projected from.
"clusterN": (AVOID) BPRMeth names clusters with numbers and numbers do not corresponds across species or TFs. This field is to be avoided
"Clusters": name of cluster
"CGI": whether the binding event overlaps with a CpG island
"MethFrag": methylome fragment the binding event overlaps with
"motCoord": coordinate of motifs within the TFBR
"motCpG": whether the motif contains a CpG
"RegRegion": regulatory region (active promoter, active enhancer or primed enhancer) overlapping with the TFBR
"DistanceTSS": distance of TFBR from the nearest TSS
"Gene": whether the TFBR is within an exon, intron or intergenic region 
"yesCTCFnames": list of TFs that cobind with the assayed binding event. [cobinding activity was assayed both considering and excluding CTCF. this column refers to the classification when it was included]
"yesCTCF":  number of TFs that cobind with the assayed binding event. [cobinding activity was assayed both considering and excluding CTCF. this column refers to the classification when it was included]
"noCTCFnames":  list of TFs that cobind with the assayed binding event. [cobinding activity was assayed both considering and excluding CTCF. this column refers to the classification when it was excluded]
"noCTCF": number of TFs that cobind with the assayed binding event. [cobinding activity was assayed both considering and excluding CTCF. this column refers to the classification when it was excluded]
